# Transcripts switched off at the stop of phloem unloading highlight the energy efficiency of sugar import in the ripening *V. vinifera* fruit

**DOI:** 10.1101/2021.01.19.427234

**Authors:** Stefania Savoi, Laurent Torregrosa, Charles Romieu

## Abstract

Transcriptomic changes at the cessation of sugar accumulation in the pericarp of *Vitis vinifera* were addressed on single berries re-synchronized according to their individual growth patterns. The net rates of water, sugars and K^+^ accumulation inferred from individual growth and solute concentration confirmed that these inflows stopped simultaneously in the ripe berry, while the small amount of malic acid remaining at this stage was still being oxidized at low rate. Re-synchronized individual berries displayed negligible variations in gene expression among triplicates. RNA-Seq studies revealed sharp reprogramming of cell wall enzymes and structural proteins at the stop of phloem unloading, associated with an 80% repression of multiple sugar transporters and aquaporins on the plasma or tonoplast membranes, with the noticeable exception of H^+^/sugar symporters, that were rather weakly and constitutively expressed. This was verified in three genotypes placed in contrasted thermo-hydric conditions. The prevalence of SWEET suggests that electrogenic transporters would play a minor role on the plasma membranes of SE/CC complex and the one of the flesh, while sucrose/H^+^ exchangers dominate on its tonoplast. *Cis*-regulatory elements present in their promoters allowed to sort these transporters in different groups, also including specific TIPs and PIPs paralogs, and cohorts of cell wall related genes. Together with simple thermodynamic considerations, these results lead to propose that H^+^/sugar exchangers at the tonoplast, associated with a considerably acidic vacuolar pH, may exhaust cytosolic sugars in the flesh and alleviate the need for supplementary energization of sugar transport at the plasma membrane.

## Introduction

During ripening, fleshy fruits shift from a repulsive to an attractive and nutritionally rewarding status, thanks to the induction of a finely orchestrated transcriptomic reprogramming in their pericarp, triggering convergent softening, colouration, accumulation of soluble sugars, and aroma development in unrelated species^1^. Fleshy fruits are divided into climacteric (e.g. tomato, apple, pear, etc.) and non-climacteric ones (e.g. grape, strawberry, citrus, etc.) depending on the occurrence of an autocatalytic emission of ethylene and a respiratory burst facilitating starch hydrolysis and cell wall degradation at the onset of ripening^2^. Grape, as the model of non-climacteric fruits, does not store any starch reserves and necessarily ripens on the vine, at the expense of a massive phloem unloading of sucrose^3^ tightly connected with malate breakdown and fruit expansion^4^. It was recently advanced that the high rate of sugar accumulation would raise an energy challenge considering the limited oxidative capacity of grapevine, which could be solved by the discharge of phloem-vectored K^+^ through the *VviK3.1* channel^5^. However, the ATP needed for sugar unloading critically depends on the respective expression and activities of SWEETs, sugar/H^+^ symporters and antiporters on the serial membrane interfaces from sieve elements to pericarp vacuoles, which remain to be elucidated even though the grapevine gene families of SWEETs, sucrose, and hexose transporters have been already described. Single berry kinetic data now argue in favour of a global sucrose/H^+^ exchange at the tonoplast^4^.

Transcriptomics approaches have become frequently employed in grapevine physiological studies focusing either on berry development or in responses to abiotic or biotic stresses. Several developmental studies targeted the incipit of berry ripening^6,7^ or the late phases of berry development^8–11^ for finding master regulators of key transitions^12^. However, albeit the clustering of RNA-Seq samples largely depended on sugar concentrations, in none of these works, there was a hint on the origin of the arrest of sugar phloem unloading in the ripening berry.

The grape berry development follows a double sigmoid curve^13^. The major events associated with each phase can be summarised as (i) a first growth phase resulting from cell division and expansion, depending on the accumulation of tartaric and malic acids as major osmotica in the green berry; (ii) a lag phase with no growth; (iii) the ripening phase initiated by berry softening, during which 1 M of glucose plus fructose accumulates at the expense of imported sucrose. During this last phase, malic acid is consumed, and the berry volume nearly doubles due to water influx. Before ripening, the xylem sap is the core source of water and minerals but phloem mass flow prevails after the onset of ripening, due to a switch from the symplasmic to the apoplastic unloading pathway^14^. One may consider that the berry is physiologically ripe when phloem unloading definitively stops, whereupon sugar concentration continues to increase only due to evaporation. It was accepted that sugar unloading might continue following growth cessation, so that excess water in phloem mass flow must be rejected by reverse xylem back-flow during late ripening^3,15,16^. However, such disconnection between sugar and water flows can be an artefact resulting from mixing asynchronous berries^4^. Difficulties emerge from the two to three weeks delay between ripening berries, which proves as long as the duration of the second growth phase itself, precluding the date of phloem arrest from being unambiguously defined on usual asynchronous samples^4,17^. Transcriptomic studies may help to find out the regulation of sugar concentration at phloem arrest, a first step for understanding the impact of climatic change on grape composition.

Attempts were made to sort berries by softening and density since the earliest transcriptomic studies^18^ to unravel key regulatory genes implicated in berry ripening. In 2008, a pioneering study^19^ investigated individual berries sorted by firmness and colour to study the early ripening process. In another work, individual berries were also considered to assess if the ripening program was accelerated in “delayed” berries^20^. Following density sorting, attempts were made to identify genes expressed or repressed at the late stages of ripening, which may be used as indicators of maturity for oenological scope^8,21^. Individual berries from vines that underwent heat stress, leaf-roll virus infection or after ozonated water applications were re-synchronized based on their primary metabolites content^22–25^. This brief survey shows that the awareness of berry heterogeneity is a rising topic, although usual samples aiming to represent the whole diversity at the experimental plot still predominate^7,26^.

In this work, we tackle the long-standing issue of sugar accumulation mechanisms in grapes upon elucidating which transcriptomic reprogramming would trigger phloem switch-off at ripe stage. Pervasive bias caused by averaging asynchronous fruits was discarded upon sorting single berries according to their actual rate of water and sugars accumulation. Prevalent transcripts stably identified in different genotypes and environments highlight the functional bases of the accumulation of 1 M hexose in this non-climacteric fruit.

## Results

### Experimental setting and monitoring of single grape berry growth during ripening

The experiment was conducted with three genotypes of *V. vinifera*: cv. Syrah, grown outdoors, and two microvines (MV032 and MV102), grown in a greenhouse, resulting in drastically different environments (Fig. S1). Although MV032 and MV102 were classified as high sugar accumulator genotypes among a segregating microvine population^27^, they showed a 35% lower sugar concentration at phloem arrest (Fig. 1, Table S1). Moreover, MV032 was ranked as a low-acid genotype while in comparison MV012 displayed +13% and +69% malic acid at the beginning of fruit ripening and at the phloem arrest (Fig. 1, Table S1).

**Figure 1.**
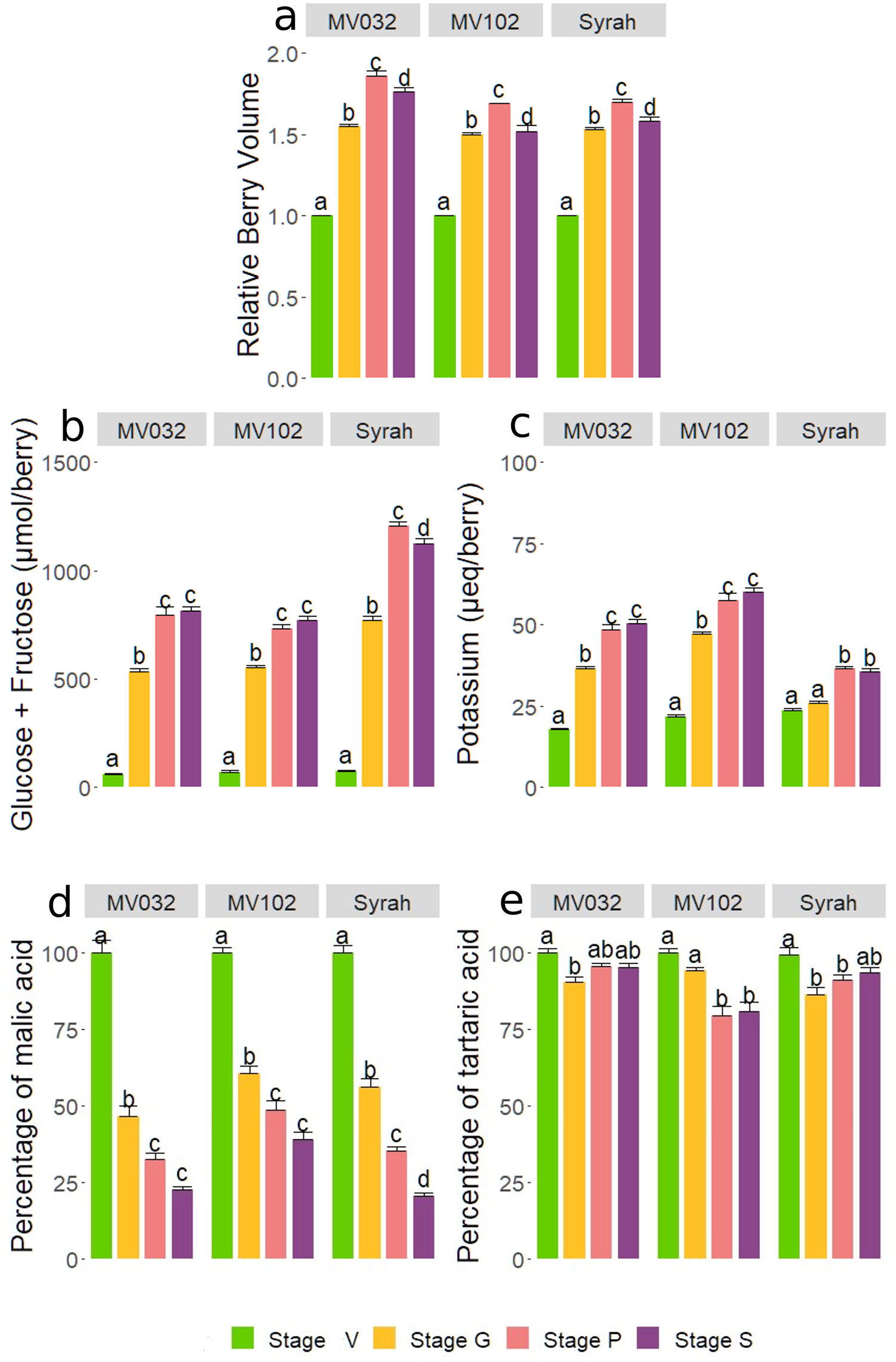
Growth and solute accumulation during the ripening of synchronised grapevine single berries of MV032, MV102, and Syrah. (a) Relative growth, (b) glucose + fructose, (c) potassium, (d) malic acid, and (e) tartaric acid. Results in b and c were normalised in order to consider 1 g of pericarp at phloem arrest. The developmental stage is indicated by colours: V for veraison in green, G for growing in yellow, P for peak in pink, S for shrivelling in purple. Letters indicate significant differences according to the Tukey ad hoc test.

Fruit growth was monitored through bi-weekly pictures and image analysis, starting from berry softening (stage V for veraison, the beginning of sugar accumulation) (Fig. 1a). Syrah, MV032, and MV102 berries were sampled at three specific stages. The first sampling, called stage G for growing, included berries growing at full speed, whose volume increased by roughly 50% within two weeks after softening. The second sampling, called stage P for peak, represented berries that were as close as possible to the peak volume, and therefore growth arrest. Lastly, the third sampling, called stage S for shrivelling, represented those berries that have peaked two weeks ago, and thus have reached the shrivelling period (Fig 1a). The changes in the total amounts of solutes per fruit (concentration x volume) confirmed that the net influx of water, photoassimilates, and K^+^ simultaneously and definitively stopped at P stage (Fig. 1, Table S1).

### Single berry primary metabolites analysis

Sugars are translocated by phloem in the form of sucrose in *V. vinifera*, and the cleaved glucose and fructose (Fig. 1b, Table S1) accumulate in equimolar amounts from stage V as the berry begins to soften. The amount of glucose + fructose in the berry dramatically increased from stage V to stage P indicating maximal activity of the apoplastic pathway of phloem unloading in G stage, accompanied by permanent cellular expansion and cell wall extension in all berries, as monitored by growth (Fig. 1a). The amount of sugars quantified in each berry did not increase any longer after the cessation of berry growth, indicating the stop of the phloem unloading. Moreover, in field conditions, the Syrah sugar content decreased in stage S, probably due to increased respiration during a heatwave recorded in those weeks (Fig. S1), while no significant decrease occurred in the greenhouse. K^+^ accumulation inside the berry (Fig. 1c, Table S1) showed the same behaviour as sugars, though its intensity was, on average, 30 and 15 times lower in Syrah and in the two microvines. K^+^ increased during the active growth phase mostly before the G stage in MV032, MV102, and after it in Syrah. There was thus no clear relationship in the timing and intensities of K^+^ and sugar unloading in the berries, except that when the phloem stopped, the amount of both solutes remained constant whatever the genotype.

Malic acid (Fig. 1d) displayed the typical pattern of malate breakdown in ripening *V. vinifera* berries, with a faster reduction between stage V and G in all genotypes. Two weeks after the cessation of phloem import (stage S), it remained between 20% (Syrah and MV032) and 40% (MV102) of the initial malic content measured in V stage. To notice that MV102 classified as high-acid genotype kept a higher amount of malic acid in shrivelling berries. It is widely accepted that tartaric acid is not metabolised during berry ripening. Indeed, in Fig. 1e, one can observe that Syrah and MV032 kept more than 90% of their initial tartaric acid content, while tartrate was reduced by less than 20% in MV102.

### Transcriptomic overview

For RNA-sequencing, nine berry pericarps per genotype (3 berries for each stage G, P, and S) were carefully selected (Fig. S2) according to their individual growth rate and primary metabolites contents. A principal component analysis (PCA) was performed over the 27 berry pericarp transcriptomes (Fig. 2a). The first and the second principal component (PC1 and PC2) accounted for 37% and 21% of the variance in gene expression among samples. Sample distribution was driven by both the developmental stage and genotype, with a clear distinction between Syrah and the two microvines. G stage samples were resolved from those at P and S stage in Syrah and MV032, while in MV102 stage distinction was less obvious. Except for one MV032 sample, stages P and S samples were mainly clustered in all genotypes. Cluster dendrogram analysis (Fig. 2b) divided the samples into four main clades. Genotypes fell in different branches that were subdivided according to the stage, except for MV102-P-2 that was closer to MV102-G-3.

**Figure 2.**
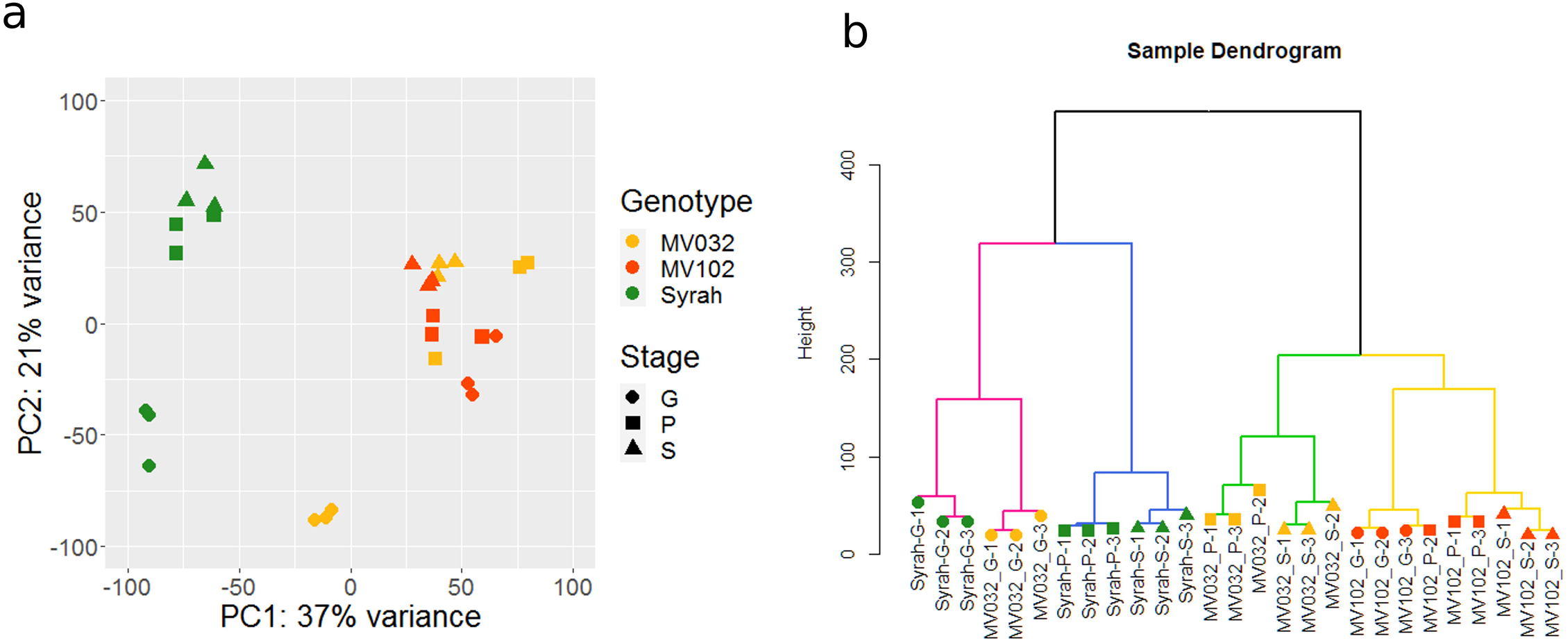
Overview of the transcriptomic results: (a) principal component analysis, and (b) sample dendrogram. Colours and symbols indicate genotypes and stages, respectively: MV032 in yellow, MV102 in red, Syrah in green; a circle for stage G, a square for P, and a triangle for S.

Overall, the sampling strategy based on image analysis and primary metabolites concentration resulted in the adequate clustering of the biological triplicates with very few exceptions found especially in the most difficult physiological stage to detect, stage P.

### Time significant genes

A comparable number of genes displayed significant (P<0.05) developmental changes in expression in the three genotypes (7025, 6629, and 6505 in Syrah, MV032, and MV102, Table S2a). They were further grouped according to their expression profiles (Fig. S3), and the cluster number was restricted to four upon visual analysis. Cluster A and B included the majority of the genes increasing or decreasing in expression from G to S (Fig. 3a). The genes commonly regulated in the three genotypes, as shown in the Venn diagram in Fig 3b, were then screened for enrichment in GO categories in the Biological Process (BP), Cellular Component (CC), and Molecular Function (MF) groups. Ultimately, clusters C and D showed genes up-peaking or down-peaking in stage P (Fig. S4a). Stage P was the most problematic stage to detect, as highlighted by the Venn diagram (Fig. S4b) where very few genes if not zero were in common between two genotypes, probably because the gene expression peak just represented small erratic variations. Growth and phloem switch-off should be a fast transition; therefore P stage detection critically relies on the accuracy of the growth measurement method and can be only characterised as *a posteriori*. The time needed to screen berries by image analysis forced us to sample the next morning when some metabolic processes were already stopped or partially modified.

**Figure 3.**
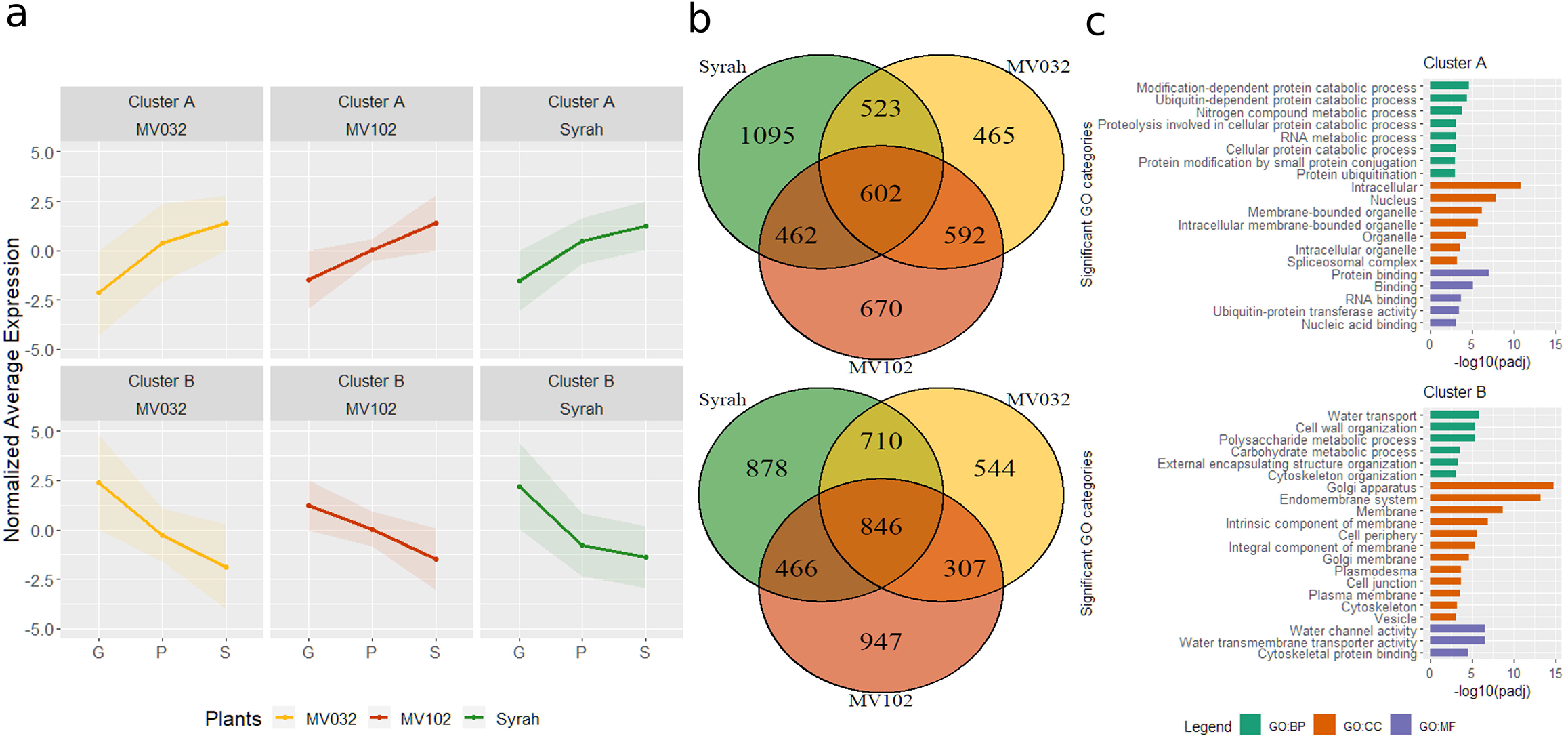
Significant time-course genes expression of (a) cluster A and B representing genes increasing or decreasing respectively in expression at phloem arrest. (b) Venn diagram showing genes displaying common or genotype specific changes in expression at phloem arrest. (c) Significantly enriched GO categories in modulated genes, divided into biological process (BP), cellular component (CC), and molecular function (MF).

Clusters A included genes whose expression increased as growth and phloem arrested and berries started to shrivel (Fig. 3a, Table S3). Six-hundred-two genes were commonly modulated in the different genotypes (Fig. 3b), whose enriched GO categories are listed in Fig. 3c. Of interest, we observed catabolic process-related categories and RNA metabolic process for BP, encoding proteins with a principal localisation in the intracellular space, but also nucleus and organelle for CC, and several binding categories were over-represented in MF. Diverse transcription factors belonging to the NAC (7 up-regulated genes), ERF/AP2 (7), bZIP (7), WRKY (7), LBD (2) multigenic families together with RNA helicases, translation initiation and splicing factor subunits and chromosome organization-related genes were up-regulated. Consistently, chromosome organisation and regulation of transcription were already reported as up-regulated in late stages of ripening^10^.

Cluster B encompassed genes decreasing in expression during development (Fig. 3a, Table S3), showing a higher number of commonly modulated genes in the three genotypes (846) (Fig. 3b). Noticeably, the BP enriched GO categories included “Water transport”, “Cell wall organisation”, “Polysaccharide and Carbohydrate metabolic process”, while for CC several categories were related to membranes, and “Water channel activity” and “Water transmembrane transporter activity”, were highly represented in MF (Fig. 3c). Among all, seven PIPs (*VviPIP1.3*, *VviPIP2.5*, *VviPIP2.3*, *VviPIP1.4*, *VviPIP2.7*, *VviPIP2.4*, *VviPIP1.2a*) and three TIPs (*VviTIP1.2*, *VviTIP1.3*, *VviTIP2.1*), sugar transporters such as *VviHT6*, *VviSWEET10*, the vacuolar invertase *VviGIN2*, the alcohol dehydrogenase *VviADH2* were highly down-regulated in the three genotypes. Besides, numerous genes related to the cuticle or the cell wall metabolism encompassing cellulose, pectin metabolism, and several expansins were strongly down-regulated. As examples, we observed *VviCER3-like*, *VviPAS2-like,* and a *VviCYP716A-like*, which are involved in the cuticular aliphatic wax biosynthesis^28^, cellulose synthases A and cellulose synthase-like, a pectate lyase, several fasciclin-like arabinogalactan proteins, pectin methyl esterases, polygalacturonases, and expansins A or expansins A-like such as *VviEXPA04*, *VviEXPA11*, *VviEXPA18*, *VviEXLA01*. Moreover, several genes related to the glycolysis/gluconeogenesis pathways, such as a UTP-glucose-1-phosphate uridyltransferase, two triose-phosphate isomerases, a glyceraldehyde-3-phosphate dehydrogenase, a phosphoglycerate mutase, one enolase were down-regulated together with malate dehydrogenases.

### Pairwise differential gene expression (S vs G)

Other important differentially expressed genes (DEGs) could be identified in Syrah and only one microvine due to discrepancies at P stage. Since the variance among P stage samples was certainly exacerbated by abrupt changes in physiology and gene expression at phloem arrest, worsened by sampling delay, data were tested in pairwise mode analysing stage S versus stage G.

A total number of 6585, 6453 and 5621 DEGs were identified in Syrah, MV032, and MV102 (Table S2b). The commonly modulated genes increased to 899 up-regulated genes and 1155 down-regulated ones, yielding to 2054 DEGs (Table S3). Other 411 genes were modulated in the three genotypes but with a discordant gene trajectory. Compared to the previous analysis, this second comparison gave a more comprehensive list of commonly modulated DEGs (before they were 1448). These two lists shared 809 genes. In this new DEGs list, there were down-regulated genes like *VviTMT2*, *VviHT2*, *VviGIN1, VviH^+^ATPase, VviEXP19* and other cell wall-related genes (Table S3).

### Rates of gene expression: ranking genes by transcription priority

Gene expression burdens cells by consuming resources and energy for transcription, translation, and the synthesis and degradation of hydrophobic proteins like transporters is particularly expensive^29^. In this respect, the transcription of transporters must stop very fast once they are no longer required. As a proxy for energy cost, we merged and ranked the two lists of differentially expressed genes according to their absolute change in expression (CPM, i.e. uncorrected for transcript length) during the stop of water, sugar and K^+^ influx in the berry (S versus G), as summarised in Table 1 (full list in Table S3). This straightforward procedure proved extremely efficient in the detection of channels and transporters virtually turned off simultaneously with phloem together with cell wall-related genes. Amazingly, in Syrah, among solute transporters, *VviHT6* exhibited the largest decay in expression from G to S stages with 80% inhibition, indicating tight synchronism with phloem arrest. This was also verified for *VviSWEET10*, *VviPIP1.3* and *VviPIP2.5*, which decreased by 90% and 83%. Moreover, a pectate lyase was virtually shut down (−98%) with several expansins (*VviEXPA19*, *VviEXPA14*, *VviEXPA16*, −73%, −92%, −99%). On the other hand, genes annotated as hydroperoxide lyase (+244%), metallothionein (+213%), and polyubiquitin (+73%) appeared as the most induced ones.

**Table 1.** List of thirty-five selected genes related to cell growth, water and sugar transport sorted by absolute difference in Syrah G versus P. Gene ID is reported as VCost.V3, V1 and RefSeq code. Gene expression is expressed as count per million (CPM).

| VCost.V3 | RefSeq | Syrah G (CPM) | Syrah P (CPM) | Syrah S (CPM) | Variation Syrah (%) | Difference Syrah | Rank Syrah | Annotation |
| --- | --- | --- | --- | --- | --- | --- | --- | --- |
| Vitvi05g01947 | LOC100264738 | 5380 | 974 | 376 | -93 | -5003.9 | 1 | early nodulin-75-like |
| Vitvi04g00442 | LOC100252479 | 4387 | 720 | 241 | -95 | -4146.8 | 2 | Uncharacterized |
| Vitvi02g00310 | LOC100233001 | 4818 | 1336 | 814 | -83 | -4004.5 | 3 | [Aquaporin] VviPIP1-3 |
| Vitvi16g01800 | LOC100232884 | 8635 | 6627 | 5693 | -34 | -2941.5 | 4 | [Cell wall - SWITCH] invertase/pectin methylesterase inhibitor |
| Vitvi13g00605 | LOC100233002 | 2773 | 971 | 462 | -83 | -2311.3 | 5 | [Aquaporin] VviPIP2-5 |
| Vitvi18g00189 | LOC100245385 | 2812 | 1299 | 751 | -73 | -2061.0 | 8 | [Cell wall] Expansin A (VvEXPA19) |
| Vitvi13g00172 | LOC100244917 | 1941 | 414 | 161 | -92 | -1780.1 | 13 | [Cell wall] Expansin A (VvEXPA14) |
| Vitvi18g00056 | LOC100232977 | 1609 | 345 | 302 | -81 | -1307.6 | 21 | [Sugar] Tonoplast Monosaccharide Transporter HT6 |
| Vitvi05g00953 | LOC100232902 | 1143 | 72 | 17 | -98 | -1125.5 | 24 | [Cell wall] Pectate lyase |
| Vitvi08g01038 | LOC100233094 | 1045 | 105 | 74 | -93 | -971.6 | 26 | [Aquaporin] VviPIP2-3 |
| Vitvi12g02137 | LOC100233113 | 1185 | 499 | 250 | -79 | -934.2 | 27 | [Cell wall] Pectin methylesterase |
| Vitvi06g01329 | LOC100265471 | 1947 | 1129 | 1070 | -45 | -876.4 | 29 | [Cell wall - SWITCH] xyloglucan endotransglucosylase/hydrolase 32 |
| Vitvi04g00465 | LOC100254408 | 1092 | 388 | 308 | -72 | -783.6 | 32 | [Cell wall] Cellulose synthase CESA1 |
| Vitvi08g01602 | LOC100233004 | 823 | 163 | 101 | -88 | -721.4 | 34 | [Aquaporin] VviTIP1-2 |
| Vitvi05g00548 | LOC100245945 | 605 | 52 | 45 | -93 | -559.8 | 40 | Sulfate transporter 3.1 |
| Vitvi15g01110 | LOC100240701 | 827 | 360 | 322 | -61 | -504.8 | 47 | [Aquaporin] VviPIP1-4 |
| Vitvi17g00070 | LOC100267921 | 511 | 71 | 48 | -91 | -462.8 | 52 | [Sugar] bidirectional sugar transporter SWEET10 |
| Vitvi15g00643 | LOC100253046 | 1015 | 413 | 554 | -45 | -460.7 | 53 | [Cell wall - SWITCH] Expansin B (VvEXPB04) |
| Vitvi03g00247 | LOC100264011 | 594 | 212 | 143 | -76 | -451.2 | 55 | [Sugar] Tonoplast Monosaccharide Transporter TMT2 |
| Vitvi18g00008 | LOC100242715 | 441 | 80 | 72 | -84 | -369.5 | 72 | [Cell wall] Cellulose synthase CESA2 |
| Vitvi03g00155 | LOC100233027 | 434 | 139 | 70 | -84 | -363.9 | 73 | [Aquaporin] VviPIP2-7 |
| Vitvi11g00272 | LOC100233140 | 541 | 259 | 177 | -67 | -363.6 | 74 | Malate dehydrogenase (NADP+) |
| Vitvi04g01668 | LOC100232854 | 1434 | 786 | 1084 | -24 | -350.1 | 78 | Alcohol dehydrogenase 2 |
| Vitvi16g00713 | LOC100256970 | 351 | 24 | 15 | -96 | -335.9 | 82 | [Sugar] Vacuolar Invertase GIN1 |
| Vitvi14g01977 | LOC100244103 | 297 | 2 | 3 | -99 | -294.2 | 93 | [Cell wall] Expansin A (VvEXPA16) |
| Vitvi18g00397 | LOC100232961 | 333 | 119 | 52 | -84 | -280.7 | 105 | [Sugar] Hexose transporter HT2 |
| Vitvi03g00209 | LOC100258952 | 286 | 35 | 25 | -91 | -260.4 | 113 | [Cell wall] Expansin A-like (VvEXLA01) |
| Vitvi18g01628 | LOC100255463 | 351 | 133 | 106 | -70 | -245.8 | 124 | [Cell wall] Cellulose synthase CESA2 |
| Vitvi11g00835 | LOC100248411 | 277 | 102 | 47 | -83 | -229.9 | 131 | [Cuticle metabolism] CER3like - Aliphatic wax |
| Vitvi12g00721 | LOC100242544 | 237 | 9 | 9 | -96 | -227.4 | 134 | fasciclin-like arabinogalactan protein 2 |
| Vitvi13g00255 | LOC100233003 | 245 | 48 | 32 | -87 | -213.6 | 143 | [Aquaporin] VviTIP1-3 |
| Vitvi13g01731 | LOC100256811 | 289 | 126 | 133 | -54 | -156.0 | 186 | Cellulose synthase CESA3 |
| Vitvi01g01805 | LOC100232906 | 157 | 7 | 3 | -98 | -153.9 | 189 | Xyloglucan endotransglucosylase/hydrolase protein B |
| Vitvi17g01251 | LOC100261426 | 136 | 7 | 2 | -99 | -134.2 | 225 | Expansin A (VvEXPA18) |
| Vitvi06g00281 | LOC100233093 | 126 | 23 | 12 | -91 | -113.9 | 281 | [Aquaporin] VviPIP2-4 |

### Promoter analysis

Sixty selected genes related to water, sugar, and growth down-regulated at the stop of phloem were distributed in different regulatory networks when clustered according to the Z-score^30^ which accounts for length, number and position of their promoter motifs (Fig. 4). C2C2dof, NAC, G2like, Homeobox, WRKY, MYB, and AP2/EREB families were driving the major differences. Some TFs displayed expression patterns consistent with the presence of target CREs in promoters of massively down-regulated genes. Interestingly, AP2/EREB and NAC binding site motifs present in *VviHT6* were mainly absent from *VviSWEET10* promoter. Among the most up-regulated NACs, we identified *VviNAC18 (Vitvi19g00271)* and *VviNAC03 (Vitvi10g00437),* previously presented as the two closest tomato NOR orthologs^31^, together with *VviNAC60* (*Vitvi08g01842*) and *VviNAC11* (*Vitvi14g01985*) putatively associated with the ripening process^31^. Conversely, WRKY motifs in *VviSWEET10* were absent from *VviHT6*, and consistently, for example, *VviWRKY47* (*Vitvi07g00523*), a new player in the sugar and ABA signaling network^32^, exhibited the largest change in expression. While MYB motifs were close to TSS in SWEETs, they were far from *VviHT6* one. Moreover, differences appeared among PIP or TIP isogenes: for example, *VviPIP2.4* and *VviPIP2.5* belonged to the same branch as SWEET while *VviPIP1.4* and *VviPIP2.3* showed a different profile. Such different regulatory networks may indicate preferential expression of these down-regulated transporter genes in specific tissues inside the pericarp, such as phloem SE/CC, hypertrophied cells in the flesh or subepidermal tissues where fruit respiration mostly occurs. Moreover, such CREs analysis was found noticeably stable in *Vitis vinifera* cultivars sequenced so far, as verified for the eleven most significant aquaporins, sugar transporters and cell wall metabolism genes discussed here, which clustering remained essentially unaffected by the genotype (Fig. S5). Such indirect approach may compensate difficulties in separating peripheral and axial phloem vessels from flesh cells turning excessively labile during ripening, while marked textural changes prevent to reach reproducible cross contaminations at different stages.

**Figure 4.**
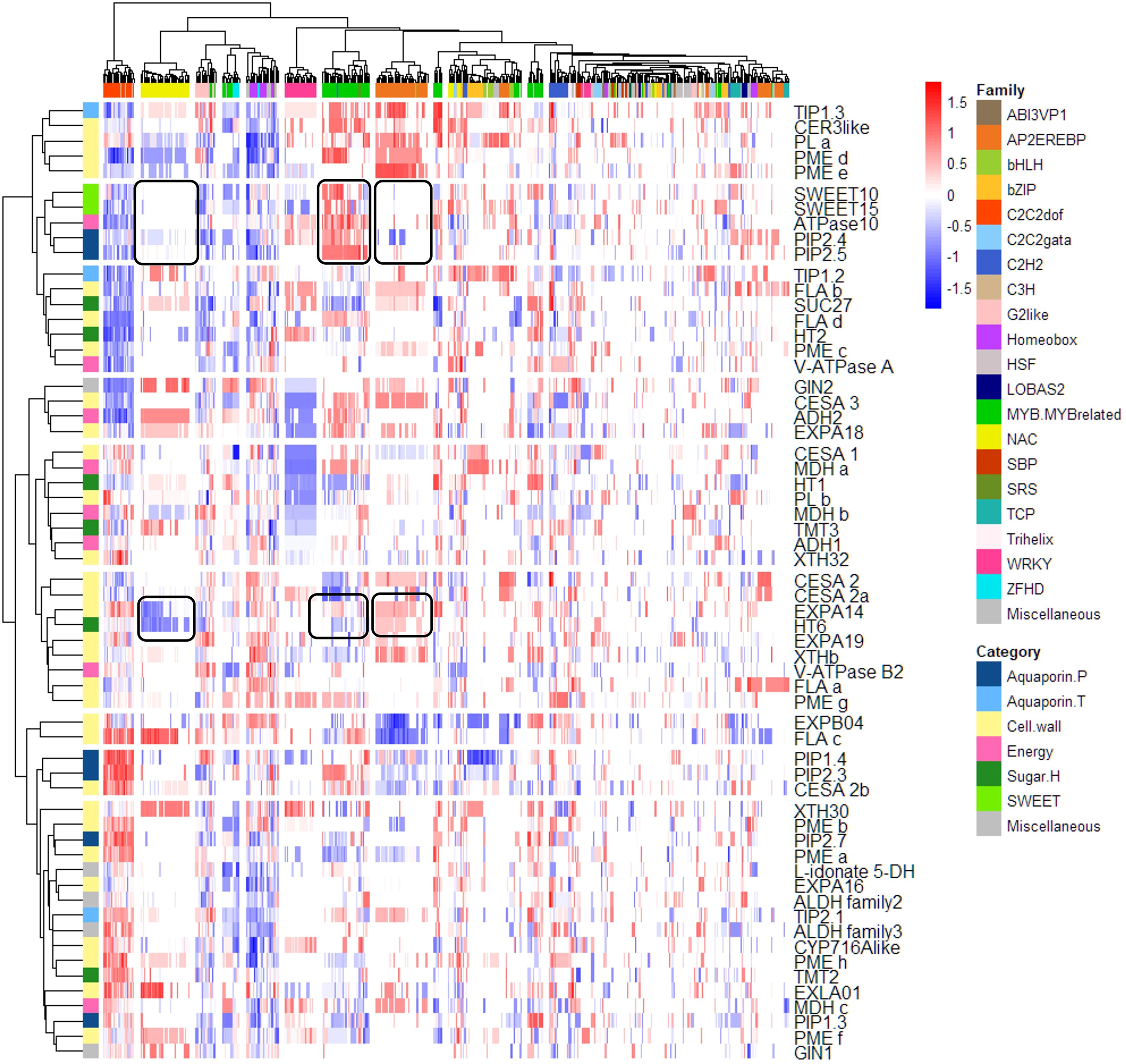
Genes inhibited at phloem stop belong to different regulatory networks. Genes and transcription factors were clustered according to the Z-score of the motifs present within 1500 bp before the TSS. Red and blue colours indicate proximity or distance of the promoter sequence to the TSS. White colour denotes the absence of a specific TF in the promoter of a gene. Horizontal and vertical colour bars grouped promoter and genes in promoter families and gene functions as indicated in the legend. The two examples discussed in the text regarding SWEETs and HT6 with reference to NAC, MYB, and AP2/EREB are marked by black squares.

## Discussion

For the first time, non-destructive monitoring of single berry growth allowed us to sample synchronised fruits within an original population marked by a significant delay in the development of individual fruits. Sorting berries according to their net growth (or water import) rate warranted that berries still growing and importing water were not mixed with shrivelling (over-ripe) ones. Real-time individual growth monitoring outperformed our previous sampling procedure which just relied on solute concentration^4,17^ upon adding the lacking kinetic dimension. Therefore, the simultaneous use of four distinct variables (malate, glucose, fructose and volume) empowered us to discriminate individual berry stages and obtain a recognizable transcriptomic signature. We showed here that the net imports of water, sugar, and K^+^ simultaneously stop when the ripening berry reaches its maximum volume, which confirms that sugar accumulation at constant volume indicating intense xylem back-flow during late ripening^3,16^ resulted from an averaging artefact on asynchronous samples^4^. This was verified in contrasted thermo-hydric conditions (field – higher temperature and water demand, versus greenhouse – non-limiting water supply) and when phloem unloading arrested at 0.8 M of sugar in microvines (as already observed for these genotypes^33^) and at 1.2 M in Syrah. At phloem arrest, with a malate concentration below 100 mEq (39 in Syrah, 48 and 93 mEq in MV032 and MV102), the berry already consumed between 60 and 80% of the malic acid accumulated before softening. These differences in acidity that would principally originate from the green stage accumulation period^27^ did not apparently interfere with phloem arrest.

Our study, performed in two environments and in three genotypes displaying different ripening features, reports for the first time robust gene trajectories associated with the end of the sugar and water accumulation processes, with the most intensively down-regulated genes related to cell wall modification (arrest of growth), aquaporins and sugar transporters (arrest of phloem unloading). The prevalence of plasma-membrane channels and transporters genes virtually turned off at phloem arrest re-emphasizes the central role of the apoplasmic pathway in ripening berry. It clearly illustrates that a strategy addressing the single berry level can lead to more comprehensive insights on fruit developmental biology than random samplings.

### The arrest of growth is modulated by the down-regulation of cell wall associated genes

The large cohort of cell wall related genes being repressed at growth arrest yields an extremely dynamic image of the rearrangements accompanying cell wall extension and cellular expansion. This cell wall life seemed to be extremely reduced after growth cessation and phloem arrest where many genes were down-regulated. The present single berry pericarp profiling greatly affirms the connection between aquaporins, cell wall and sugar transport. No obvious marker of cell death was detected here, which is often described later in the shrivelling process of Syrah berries^34^.

Here we reported that genes involved in cellulose metabolism (15 cellulose synthases), hemicellulose metabolism (4 endo-1,4-beta-glucanases, 7 xyloglucan endo-transglucosylase / hydrolase proteins), pectin metabolism (4 pectate lyase, 4 polygalacturonases, 6 ß-galactosidases, 6 fasciclin-like arabinogalactan protein), and expansins genes (9) were strongly down-regulated at the arrest of phloem (Table S3). In particular, we emphasise that precisely such expansins (*VviEXPA19*, *VviEXLA1*, *VviEXPB4*, *VviEXPA14*) previously linked to growth and cellular expansion during the second growth phases both in the flesh and in the skin^35^ were down-regulated concomitantly with growth cessation. A link between these expansins and aquaporins has been envisaged in a pre-genomic study reporting a down-regulating trend two or three weeks before the attainment of maximum berry size on asynchronous berry samples^36^. This so-called “unexpected” link was recently confirmed through a gene coexpression network analysis^37^.

### The arrest of phloem is linked with strongly down-regulated aquaporins

Aquaporins are transmembrane channels facilitating the transport of water and small solutes from cell to cell and between cell compartments. Noticeably, seven of the ten plasma membrane intrinsic proteins (PIPs) identified in *V. vinifera*^37^ were drastically inhibited (Table 1, Table S3) simultaneously with growth and phloem arrest. Among them, *VviPIP1.3* (*Vitvi02g00310 - LOC100233001*), *VviPIP2.5* (*Vitvi13g00605 - LOC100233002*), and *VviPIP2.3* (*Vitvi08g01038 - LOC100233094*) were the most intensively repressed transcripts (ranked 3^rd^, 5^th^, and 26^th^ according to their absolute expression changes in Syrah) with a percentage of inhibition ranging from 83 to 93% in Syrah. Data on grapevine aquaporins^37^, including the gene expression atlas^26^ and various other RNA-Seq datasets, showed that these three genes exhibited quite a common expression pattern and actually decreased at ripe or harvest stages (defined as 20 °Brix) but the inhibition continued in post-harvest withering phases (see figure 3a,b in^37^). Present results showed that this progressive decrease does not occur in individual fruits, but may represent an emerging property on asynchronous samples. While the *PIP1* gene family was more associated with dynamic changes in membrane permeability, *PIP2* was recently allocated to sieve element plasma membrane in poplar, a symplastic loader^38^. Much work remains to be done regarding their exact localisation in an apoplastic unloader such as grapevine berry insofar direct orthologous relationship between *Vitis*, *Populus*, and *Arabidopsi*s are difficult to establish^37^. Finally, among the eleven tonoplast intrinsic proteins (TIPs), only three were significantly inhibited. The TIPs expression level was lower than in the three previously discussed PIPs. In particular, the predominant *VviTIP1.2* (*Vitvi08g01602 - LOC100233004*) was six times less expressed than *VviPIP1.3* but exhibited a huge inhibition upon phloem arrest (88%).

### Phloem unloading is stopped by the down-regulation of specific sugar transporters

Phloem unloading via the apoplastic pathway involves sugar transport through sequential interfaces from the sieve element-companion cell complex (SE/CC) to sink tissues such as the hypertrophied cells of the pericarp till its final storage compartment, the vacuole.

Sugar Will Eventually be Exported Transporters (SWEETs) are sugar uniporter proteins recently identified in plants. We show here that *Vitvi17g00070* (*LOC100267921*) was the only SWEET gene highly down-regulated at the arrest of phloem. This gene, annotated as SWEET14 at NCBI, corresponds to *VviSWEET10* (*VIT_17s0000g00830*) in ^39^, the ortholog of *AtSWEET10*, a plasma membrane sucrose transporter of the clade III SWEETs. These authors^39^ evidenced a strong induction of *VviSWEET10* at veraison, without berry sorting nor compositional data, and confirmed its expression at the plasma membrane. Ectopic expression in tomato increased its sugar content, while promoter-GUS fusions showed preferential expression in the vascular bundles and the flesh. These authors privileged a *VviSWEET10* role in hexose loading in the flesh, but it may preferentially export sucrose from SE/CC to the pericarp apoplast as reported for *AtSWEET11* and *AtSWEET12*, two other *Arabidopsis* genes belonging to the same clade III SWEETs^40^. Noticeably, preferential expression of *VviSWEET10* was confirmed during ripening in Riesling and Cabernet Sauvignon with a decrement at 100-110 days after veraison^41^. However, the same authors evidenced a higher expression of *VviSWEET15 (Vitvi01g01719)* in Petit Manseng, which was not followed by a marked decrease at phloem arrest^41^. Our RNA-seq data show that *VviSWEET15* displays a similar expression level than *VviSWEET10* during the active phloem unloading without being inhibited at phloem stop (Table S4) as reported in ^39^. This might suggest that *VviSWEET10* would be preferentially expressed at the phloem unloading site while *VviSWEET15* would facilitate sugar transfer between the core and the periphery of the fruit, where respiration occurs (Fig. 5).

**Figure 5.**
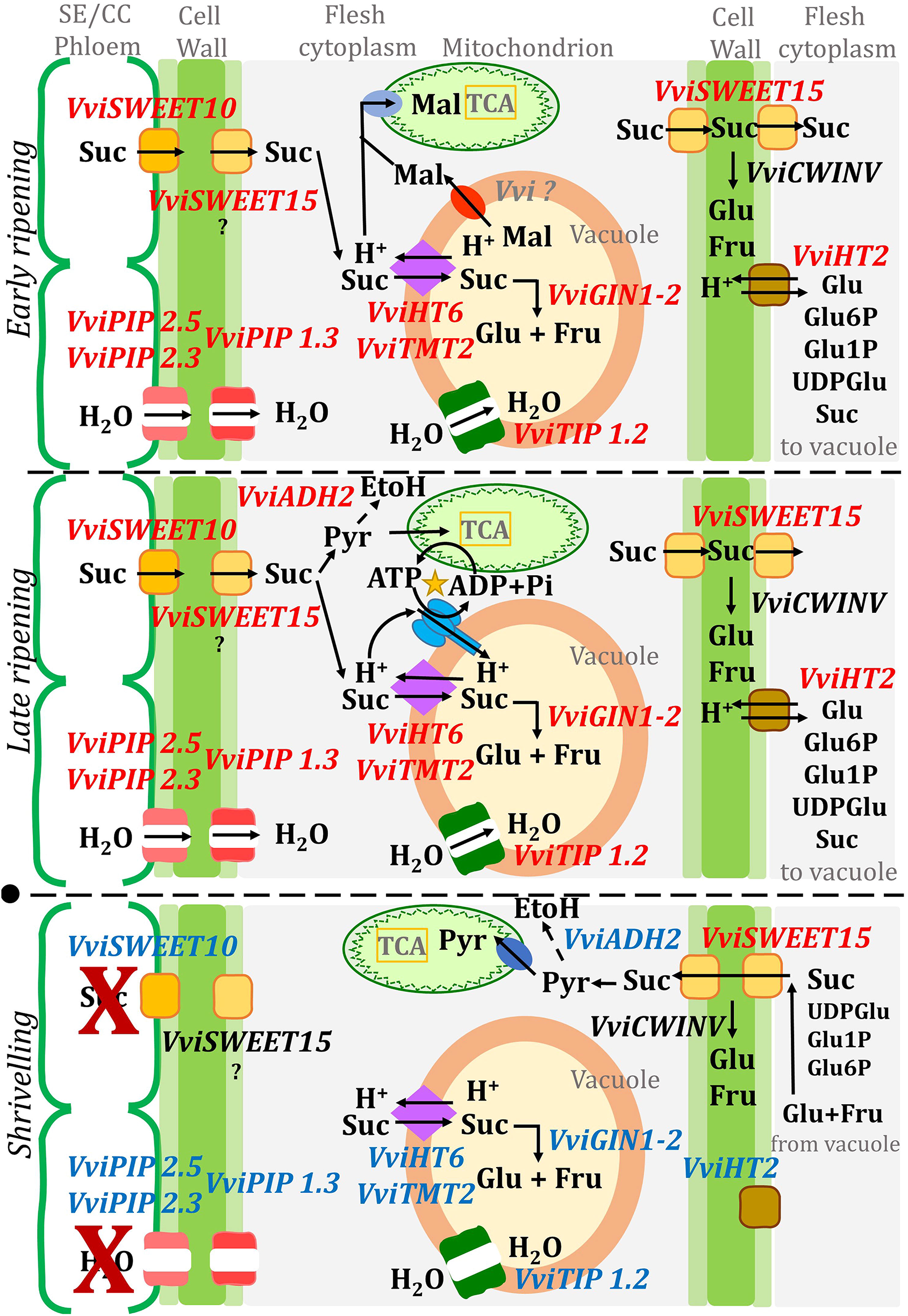
Hypothetical structure of the energy-efficient apoplastic pathway in the ripening berry. The figure is divided into three panels: early ripening, late ripening, and shrivelling with the stop of phloem between the two latter (marked by a black circle). Genes written in red or blue italics refer to those respectively expressed or repressed at specific stages, those in black are constitutive.

Conversely to SWEET, sugar/H^+^ antiporters and symporters can transport sugar against its concentration gradient using the proton motive force. *VviHT6* (*Vitvi18g00056* - *LOC100232977*) was the sugar transporter with the highest decrement in expression at the stop of phloem. *VviHT6* induction during Syrah ripening^42^ was reported before the release of the *Vitis* genome^43^ together with its complete ORF (DQ017393.1), as confirmed in other cultivars^44^. The prevalence of *VviHT6* expression over those of *VviSUT*s and *VviHT*s was pointed out^45^, but its synchronism with sugar accumulation remained particularly elusive due to excessive sampling period and absence of sugar measurements. The expected targeting of *VviHT6* to the tonoplast membrane and its accumulation during ripening were masterly confirmed through proteomic analysis of purified vacuoles^46^, while present ectopic expression of *VviHT6::GFP* fusion in the hairy root homologous system rules out its presence on grapevine plasma membrane (Fig. S6). Finally, the recent re-evaluation of malate and sugar fluxes in synchronised berries provided the first quantitative evidence that an overall H^+^/sucrose exchange parallels the predominant expression of *VviHT6* in ripening berries^4^. In line with accumulating evidence that *VviHT6* orthologs should be renamed tonoplast sugar transporter (TST1/2) in *Arabidopsis*, apple, taproot, and melon^47–50^, this important result enlights the long-standing question of the origin of the malate/sugar relation in grapevine berry. *VviTMT2* (*Vitvi03g00247* - *LOC100264011*), an isogene three times less expressed than *VviHT6* at G stage, decreased by 76, 74 and 54% in Syrah, MV032, and MV102; however, at the protein level, *VviTMT2* and *VviHT6* display comparable expression^46^. Moreover, the repression of *VviTMT2* was recently pointed out in low-sugar accumulating berries affected by the berry shrivel ripening physiological disorder^51^. The third gene of this clade, *VviTMT3* (*Vitvi07g01898* - *LOC100243856*)^44,52^ was not expressed although it was the first one localised at the tonoplast^52^.

The probable PM H^+^ hexose symporter *VviHT2* (Vitvi18g00397 - *LOC100232961*)^44^ also exhibited a considerable increase during ripening, followed by a 35% decrease in two weeks^53^. Synchronised berries show here that more than 50% inhibition occurs at growth arrest, reaching after that 84% decrease in Syrah. *VviHT2* was already associated with the induction of *VviHT6* at veraison both in flesh and skin^54^ and in ABA, and GA_3_ treated berry^45,55^, but data were lacking regarding its inhibition at ripe stage. To the best of our knowledge, *VviHT2* localisation is still unknown. Unfortunately, more information is available for *VviHT1,* but it was not commonly modulated in the genotypes studied here.

### An energetically optimised sugar accumulation pathway operates in berries?

A ripening berry can just produce 3-6 ATP by oxidative phosphorylation while it accumulates one hexose^4^. Thus, a phloem unloading pathway passing through H^+^-coupled sugar transporters on the three membrane interfaces from SE/CC complex to flesh vacuoles, combined with the functioning of apoplasmic invertase, would waste within 50% of cellular ATP for H^+^ recycling, in a pure loss, as the sucrose gradient between first and terminal compartments is thermodynamically favourable in *V. vinifera.* Present results (DEGs, promoter analysis) lead to propose a more efficient design of sugar import, which refines the previous ones^4,56^ upon integrating key genes/functions observed in this study (Fig. 5).

*VviSWEET10*, located on the SE/CC plasma membrane, would be the major player in the export of sucrose from the phloem into flesh apoplasm. Sucrose hydrolysis in the apoplasm would require its resynthesis in flesh cells to allow sucrose vacuolar transport by TST (see discussion above) which could be compatible with the huge induction of SPS during ripening^23,56^ and with SPS, SuSy, AI, NI *in vitro* activities^41^ in five thousand excess, at least, with sugar accumulation rate. However, except for AI, the corresponding genes were not strongly inhibited at phloem arrest, which may indicate that sucrose resynthesis could be also involved in intercellular sugar exchanges needed for respiration in fruit periphery, which continues after phloem arrest. In fact, *Vitvi07g00353* and *Vitvi11g00542* annotated as sucrose synthase and sucrose phosphate synthase (Table S4) were rather constant during the three stages in the three genotypes.

Sugar transport by SWEETs is energetically silent and the sink strength for sugar accumulation is driven by the key electrogenic antiporters (proton-coupled, i.e. energy consuming) *VviHT6* and *VviTMT2* that transport sucrose across the tonoplast. At the beginning of ripening, the vacuolar acidity within 400 and 450 mEq (Table S1) at pH 2.7, provides a considerable proton motive force for the transport of sucrose into the vacuole, as long as the counterion malate is available. In tandem with vacuolar invertase, this system is susceptible to restrict cytosolic sucrose below the nanomolar range. Although less dramatic, a similar conclusion holds for hexose, in case *VviHT6* and *VviTMT2* do not transport sucrose (Table S5). Once in the cytoplasm, the exchanged malic acid will serve as a substrate for respiration or gluconeogenesis, according to the reactions: 2 H^+^ + malate + 3 O_2_ = 4 CO_2_ + 3 H_2_O and 2 malate + 4 H^+^ → 1 hexose + 2 CO_2_ that progressively vanish with the exhaustion of malic acid. Between 22 (Syrah, exposed to stress) and 40% (MV032, greenhouse) of the H^+^ exchanged with sucrose at the tonoplast (Table S1) are scavenged from the cytoplasm through these reactions, but the remaining ones need to be pumped back in the vacuole at the expense of a progressive activation of glycolysis, aerobic fermentation (ADH2), vacuolar H^+^/ATPase and H^+^/PPiase^57^. It is noticeable that also *VviADH2* is inhibited at phloem arrest, along with vacuolar invertase. Results on the respiration of single grapevine berry, which provide strong arguments in this direction, will be detailed in a next publication. The only PM H^+^ symporter encountered here (*VviHT2*) displayed the highest expression in Syrah, which also needed more ATP at the tonoplast (Table S1), when compared to microvines. This suggests that energy-requiring H^+^ symporters would be activated on the plasma membrane to increase sink strength in unfavourable environmental conditions, while *VviSWEET15* would allow the bulk of sugar import on flesh cells, even though further work is needed to precise its substrate specificity.

The energetically-optimised model proposed here alleviates the opening of an AKT2-like channel that would compensate the energy deficit during phloem unloading^5^, fully inherent to the sugar/H^+^ symporters dogma, that can be discussed on thermodynamic and transcriptional grounds in grape berry. Such a mechanism would require a simultaneous activation of quantitatively similar flows of sugars and K^+^ in ripening berries, which must be clearly excluded now. In the absence of growth measurements, previously published correlations confused accumulation and concentration mechanisms, moreover a concomitant concentration of sugar and K^+^ occurs without any need of co-transport mechanisms in shrivelling berries after the stop of phloem. In future studies, it would be interesting to understand which aspects of the energetically optimised mechanism proposed here are conserved among cultivated and ancestral cultivars or if a genetic diversity prevails, for example in extreme cultivars such as acidless mutants or acidic varieties.

## Materials and methods

### Plants

Three genotypes were characterised in this study: the cv. *V. vinifera* Syrah, a traditional variety adapted to warm climates and used for wine production, and two hermaphroditic microvines offspring, named MV032 and MV102. Both microvines displayed a semi-dwarf stature^33^, resulting from five pseudo backcrosses between *Muscadinia rotundifolia* G52 and *V. vinifera* genotypes. The last cross involved the 04c023V0003 female microvine^58^ and the 3197-81B hybrid^59^. Genetically, microvines used here can be considered very close to *V. vinifera* (1% or less of *M. rotundifolia* background). Syrah vines grafted on SO4 were in an open field at the experimental vineyard of Montpellier Supagro planted in 2000 while the two-years-old microvines were grown in pots in a greenhouse under semi-controlled condition (25/15°C day/night temperature, VPD 1 kPa, photoperiod 12 h).

### Single berry sampling

Single berry growth and development were monitored through biweekly pictures of selected clusters (up to 8 for each genotype) starting a few days before softening day until two weeks after maximum berry growth. Pictures were taken using a Lumix FZ100 camera (Panasonic), keeping the focal range and cluster to camera lens distance (30cm) constant. The volume increment of selected berries was calculated by analysing the pictures with ImageJ^60^. The software, after calibrating the images using the 1 cm scale present in each of them, automatically counted the pixels enclosed in each targeted berry area, measured as an ellipsoid. The estimated berry volume was then calculated accordingly using the radius of the previously calculated area. To eliminate the intra- and inter-cultivars variability in berry sizes and compare the changes in volume among berries of different clusters, vines, and genotypes, each berry growth profile was normalised to the softening volume, set to 1.

According to their own volume changes, 10 to 15 berries were sampled for each genotype at different dates during the ripening growing phase (stage G), closest to the berry growth peak (stage P) and two weeks after the maximum growth (stage S) in the shrivelling phase. Concerning stage P, one must notice that *a-posteriori* it was found that phloem and, obviously, growth, were already blocked at this stage, which should, technically speaking, be called P+1. Berries were sampled at the same time of the day, between 9 am to 11 am, to avoid circadian cycle influences. Berries without pedicel were rapidly deseeded before freezing (1-2 min after harvest) in liquid N_2_ and stored at −80 until further analysis. Single berries were ground to a fine powder under liquid nitrogen using a ball mixer mill (Retsch MM400).

### Primary metabolites analysis

Glucose, fructose, malate and tartrate were analysed by HPLC. For each berry, 100 mg of powder was weighed, diluted 5x with a solution of HCl 0.25 N, well shacked, and left overnight at room temperature. Samples were then centrifuged for 10 min at maximum speed, and a supernatant aliquot was diluted 10x with a solution of H_2_SO_4_ containing as internal standard 600 µM of acetic acid. Samples were injected according to ^23^. For K^+^ analysis, the previous samples in HCl were further diluted 10x with MilliQ water and analysed as in ^27^. Tukey.HSD post-hoc test was used as a statistical method to find significant differences among metabolites concentration over time.

### RNA extraction and sequencing

Based on growth increment, internal sugars and organic acids, three berries (pericarp) per stage G, P, and S for each genotype were selected for individual RNA extraction and library preparation, as in ^23^. Samples were sequenced on an Illumina HiSeq3000 in paired-end mode, 2×150 bp reads, at the Genotoul platform of INRAe-Toulouse.

### Data analysis

Raw reads were trimmed for quality and length with Trimmomatic, version 0.38^61^. Reads were aligned against the reference grapevine genome PN40024 12×2^62^, using the software Hisat2, version 2.1.0^63^ yielding an average of 31.6 M sequence per sample (Table S6). Aligned reads were counted using the last available annotation VCost.v3 with HTSeq-count (version 0.9.1) with the “nonunique all” flag^64^. Genes were filtered by applying an RPKM>1 cut-off in at least one experimental condition (Table S4), and the variance stabilising transformation was applied for data visualisation.

Genes (TMM normalised) were tested for multi time-series significance for finding genes with significant (P<0.05) temporal expression changes using the R package MaSigPro^65^, with Syrah dataset used as a control. A quadratic regression model (rqs=0.7) was applied. Significant genes over time were clustered using STEM (short time-series expression miner) software^66^ with the maximum number of model profiles set to 24. Filtering or normalisation was not applied by the software since input data were already filtered and normalised, but a mean-centred scale was applied to overcome cluster mismatches due to different expression levels. Genes showing the same profiles pattern among the significant clusters were grouped as follows: cluster A = cluster 7 & 9; cluster B = cluster 16, 17 & 14; cluster C = cluster 8; cluster D = cluster 15, (Fig. S3). Venn diagrams were drawn with the R package VennDiagram. GO term profiler was retrieved from the web tool gProfiler by applying a significant p-value of 0.001.

Pairwise differential expressed genes (FDR◻<◻0.05) analysis was performed with the R package DeSeq2^67^ testing S/G samples. As before, genes were filtered by RPKM>1.

### In silico functional CREs analysis

Promoter sequences (1.5 kb upstream of TSS) were extracted using bedtools from the 12X2 genome assembly^62^. Gene correspondence file at https://urgi.versailles.inra.fr/content/download/5723/43038/file/list_genes_vitis_correspondencesV3_1.xlsx and in the related gff3 shows that the 5’UTR was frequently lost in the VCost.v3 annotation. Therefore, we decided to take the longest sequence between VCost.v3 and NCBI (Vitvi and LOC references, respectively) as the most probable TSS. *Cis*-regulatory elements (CREs) from a selected list of genes involved in the stop of the phloem were searched for an exact match based on the DAP-Seq motif library^68^ with HOMER^69^. For each gene, each CRE was normalised for positional bias score by calculating the Z-score, which considers the motifs length, its frequency and position^30^. To further validate the CREs results obtained from PN40024, the sequences of a subgroup of eleven genes (*PIP1.3*, *PIP2.4*, *PIP2.5*, *TIP1.2*, *EXP14*, *EXP19*, *HT6*, *HT2*, *TMT2*, *SWEET10*, and *SWEET15*) were retrieved from the genomes of Cabernet Sauvignon, Merlot, Zinfandel, and Chardonnay (http://www.grapegenomics.com)^70–73^, and their promoters were analyzed as before.

## Data Availability

All raw transcriptomics reads have been deposited in NCBI Sequence Read Archive (http://www.ncbi.nlm.nih.gov/sra). The BioProject ID is PRJNA673575.

## Acknowledgements

We would like to thank the Poupelain Foundation, the Comité Interprofessionnel des Vins de Bordeaux (CIVB) and the Agence Nationale de la Recherche (G2WAS project, ANR-19-CE20-0024) for providing the financial support of this study. Thanks to Marc Farnos for plant management, Sylvain Santoni and Muriel Latreille for RNA library preparation, Gauthier Sarah for bioinformatics support and Yannick Sire for cation analysis.

## Contributions

SS monitored and collected the grape samples, extracted the RNA and the metabolites, performed the data analysis, interpreted the results, and drafted the manuscript. LT provided the microvine lines and supervised the experiments, interpreted the results, and edited the manuscript. CR conceived and designed the study, coordinated and supervised the experiments, interpreted the results, and drafted the manuscript. All authors read and approved the final version of the manuscript.

## Supplementary information

**Figure S1.** Meteorological data showing temperature max and min recorded in the greenhouse (dotted lines) and in the field (solid line) during the grape growing season. Syrah sampling dates (V, G, P, and S) based on Julian days are reported on the top.

**Figure S2.** Glucose plus fructose (mM) in relation to malic acid (mEq) in Syrah, MV032, and MV102. Different shapes denote different developmental stages: circle, square and triangle represent stage G, P, and S. Coloured points in red, yellow and green represent the samples selected for RNA-Seq analysis in G, P and S.

**Figure S3.** Overview of the model profiles interface (from the STEM software) where each box corresponds to a model expression profile, manually set to 24. Red lines show genes distribution. Coloured profiles have a significant number of genes assigned and therefore were considered for the following analysis.

**Figure S4.** Significant time-course genes expression of clusters C and D representing genes up or down peaking respectively in expression at phloem arrest and Venn diagram showing genes displaying common or cultivar specific changes in expression at phloem arrest.

**Figure S5.** Promoter analysis of selected genes (PIP1.3, PIP2.4, PIP2.5, TIP1.2, EXP14, EXP19, HT6, HT2, TMT2, SWEET10, SWEET15) from PN40024, Cabernet Sauvignon, Merlot, Zinfandel, and Chardonnay. Gene were clustered according to the Z-score of the motifs present within 1500 bp before the TSS. Red and blue colours indicate proximity or distance of the promoter sequence to the TSS. White colour denotes the absence of a specific TF in the promoter of a gene. Horizontal and vertical colour bars grouped promoter and genes in promoter families and gene functions as indicated in the legend.

**Figure S6.** Subcellular localization of *VviHT6* proteins to the tonoplast after stable expression in grapevine hairy roots. The complete ORF of the *VviHT6* gene without the stop codon was amplified from Syrah cDNA (ripening berry) using the following primer pair: HT6for CACCATGAACGGAGCTGTGCT, HT6rev GTCATTCTTTGCTGCAGTAAC before blunt end topo cloning, transfer to the pH7FWG2 vector and stable transformation of grapevine with agrobacterium rhyzogenes A4 according to Gomez et al., 2009.

**Table S1.** Evolution of sugars, acid and cations. (a) Amounts per fruit (concentration x volume); (b) concentrations and (c) net H^+^/sucrose exchange escaping V-ATPase at the tonoplast.

**Table S2.**List of genes displaying time significant expression (P<0.05) in the three genotypes. (a) Time significant gene identified with MaSigPro (G-P-S); (b) pairwise significant gene identified with DeSeq2 (S/G).

**Table S3.** Merged list of genes showing a co-expressed pattern ranked by absolute difference in CPM expression between G and S.

**Table S4.** Genes that passed the filtering threshold (RPKM>1) in at least one group of sample triplicates.

**Table S5.** Depletion of cytosolic sugar at equilibrium state of vacuolar transports, in fruit parenchymal cells from the onset to the arrest of sugar loading.

**Table S6.** RNA-Sequencing data metrics.

