## Supplementary Figure S1-S6 for "Transcripts switched off at the stop of phloem unloading highlight the energy efficiency of sugar import in the ripening *V. vinifera* fruit"

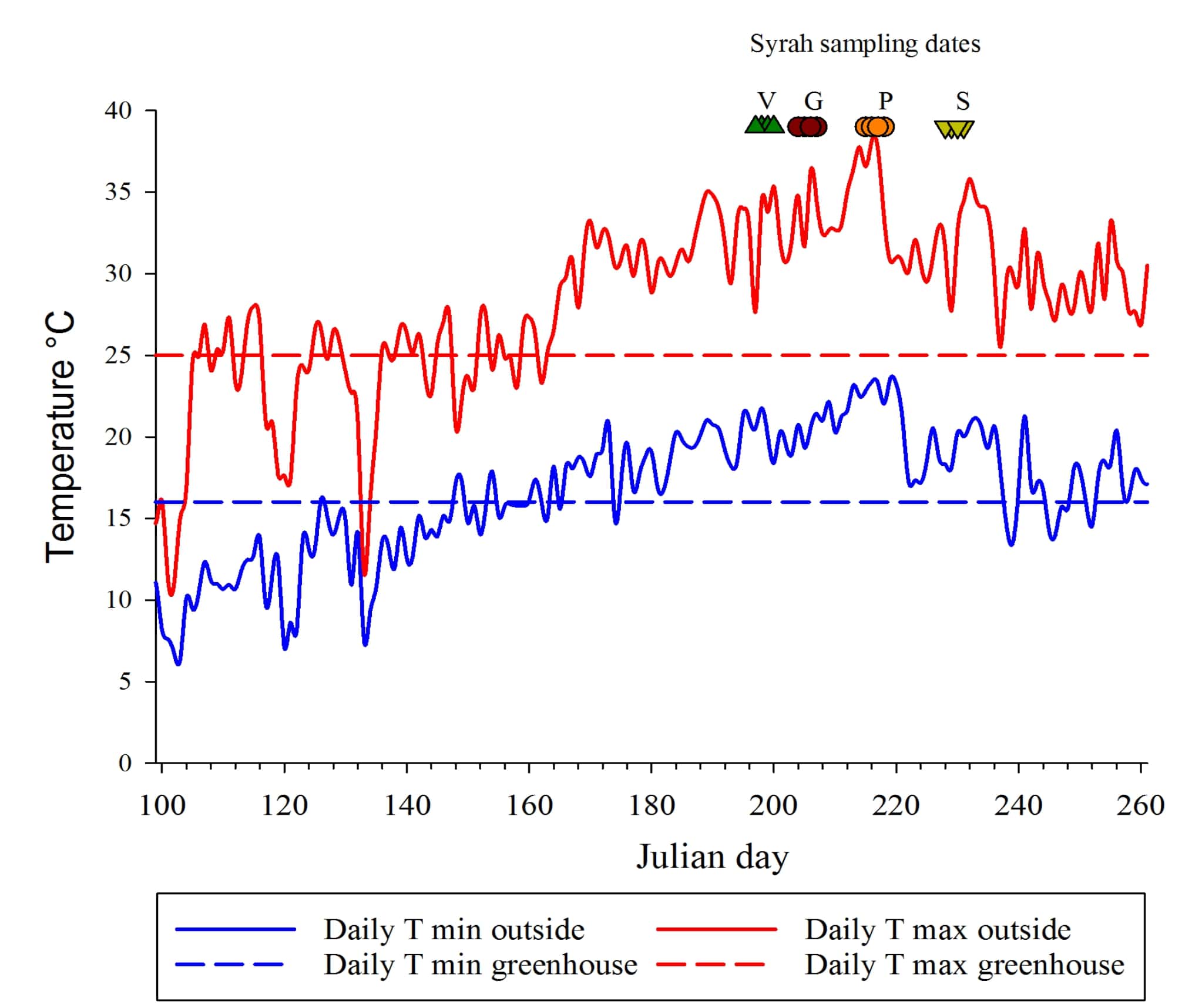

**Figure S1.**

Meteorological data showing temperature max and min recorded in the greenhouse (dotted lines) and in the field (solid line) during the grape growing season. Syrah sampling dates (V, G, P, and S) based on Julian days are reported on the top.

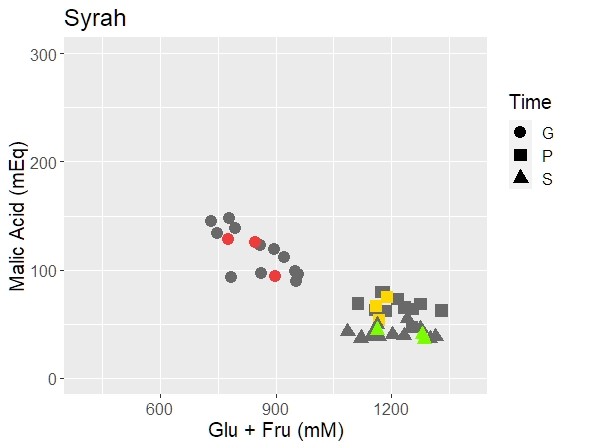

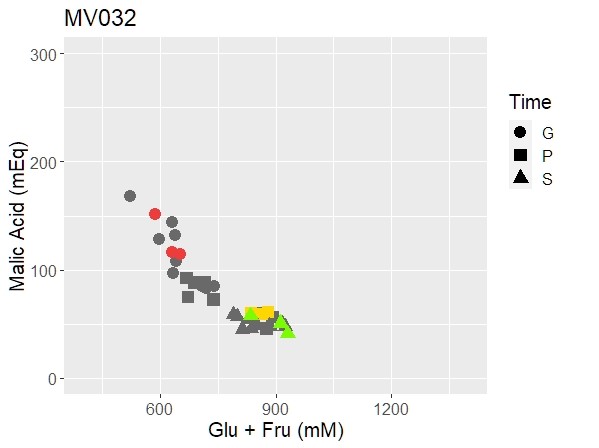

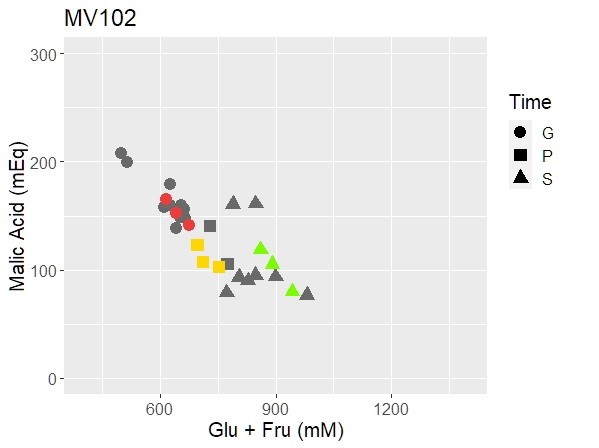

**Figure S2.**

Glucose plus fructose (mM) in relation to malic acid (mEq) in Syrah, MV032, and MV102. Different shapes denote different developmental stages: circle, square and triangle represent stage G, P, and S. Coloured points in red, yellow and green represent the samples selected for RNA-Seq analysis in G, P, and S.

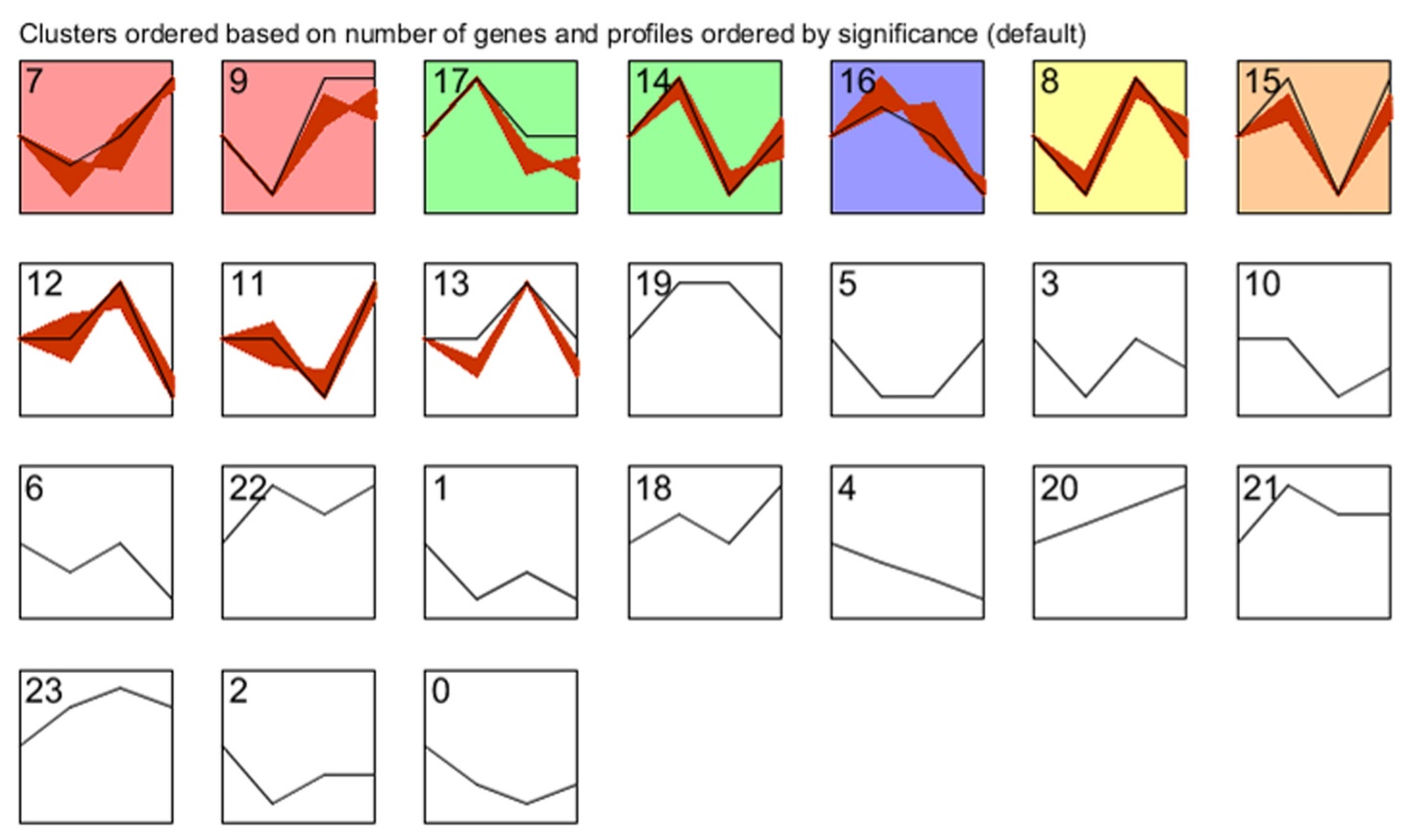

**Figure S3.**

Overview of the model profiles interface (from the STEM software) where each box corresponds to a model expression profile, manually set to 24. Red lines show genes distribution. Coloured profiles have a significant number of genes assigned and therefore were considered for the following analysis.

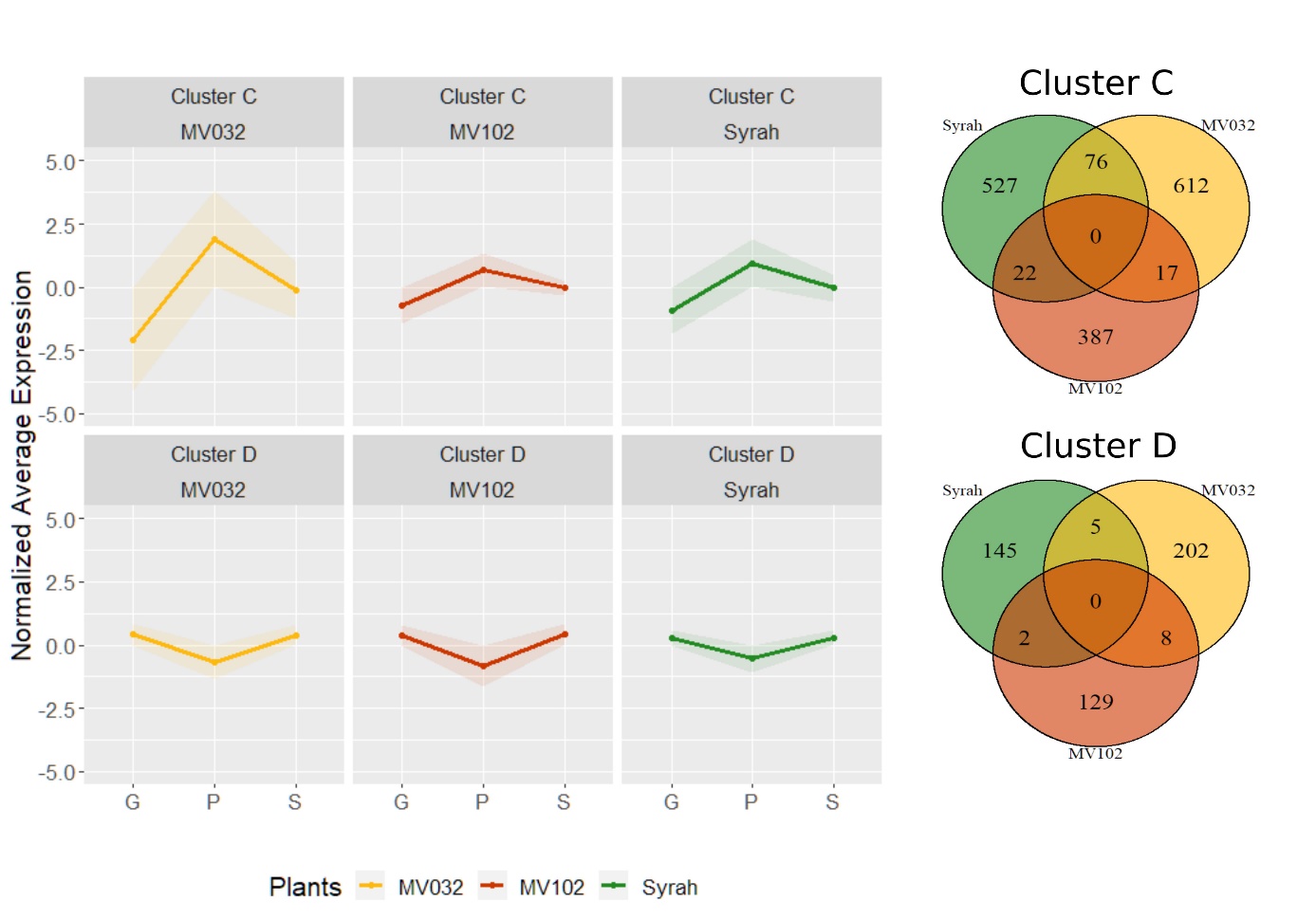

**Figure S4.**

Significant time-course genes expression of clusters C and D representing genes up or down peaking respectively in expression at phloem arrest and Venn diagram showing genes displaying common or cultivar specific changes in expression at phloem arrest.

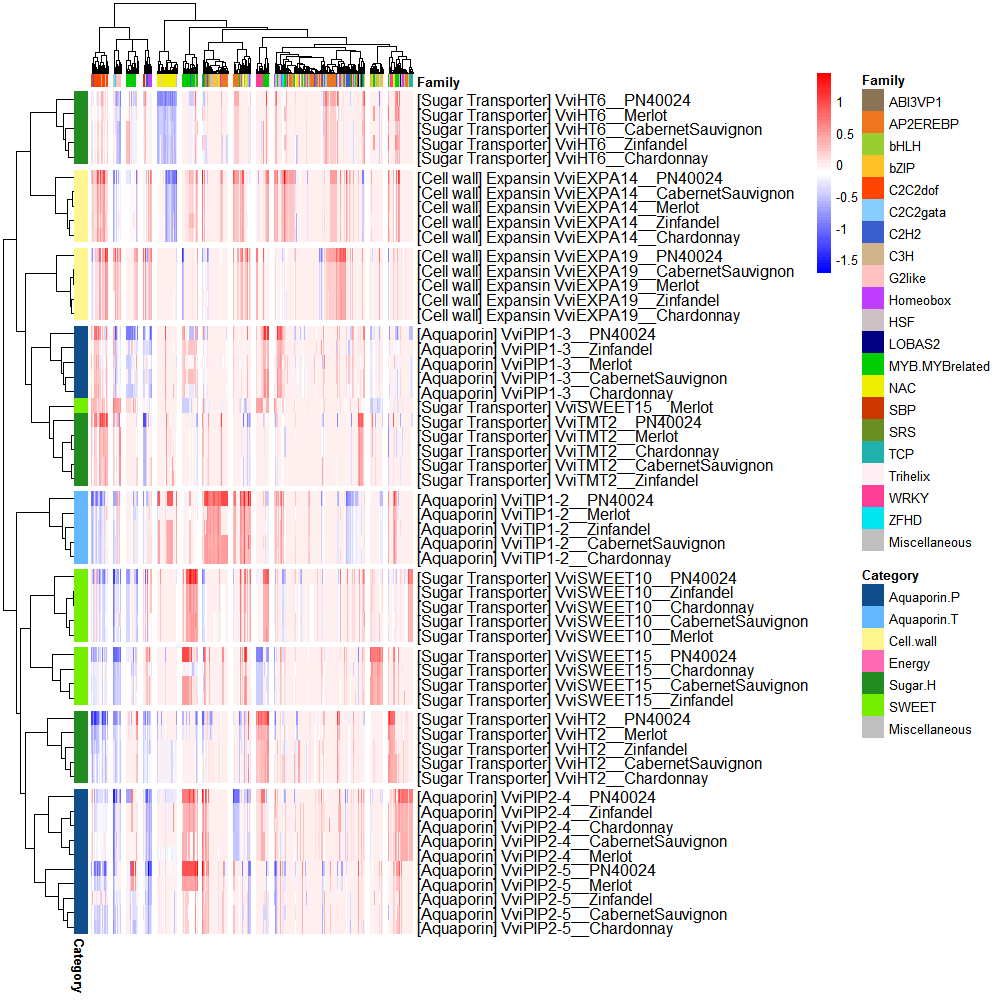

**Figure S5**

Promoter analysis of selected eleven genes (PIP1.3, PIP2.4, PIP2.5, TIP1.2, EXP14, EXP19, HT6, HT2, TMT2, SWEET10, SWEET15) from PN40024, Cabernet Sauvignon, Merlot, Zinfandel, and Chardonnay. Gene were clustered according to the Z-score of the motifs present within 1500 bp before the TSS. Red and blue colours indicate proximity or distance of the promoter sequence to the TSS. White colour denotes the absence of a specific TF in the promoter of a gene. Horizontal and vertical colour bars grouped promoter and genes in promoter families and gene functions as indicated in the legend.

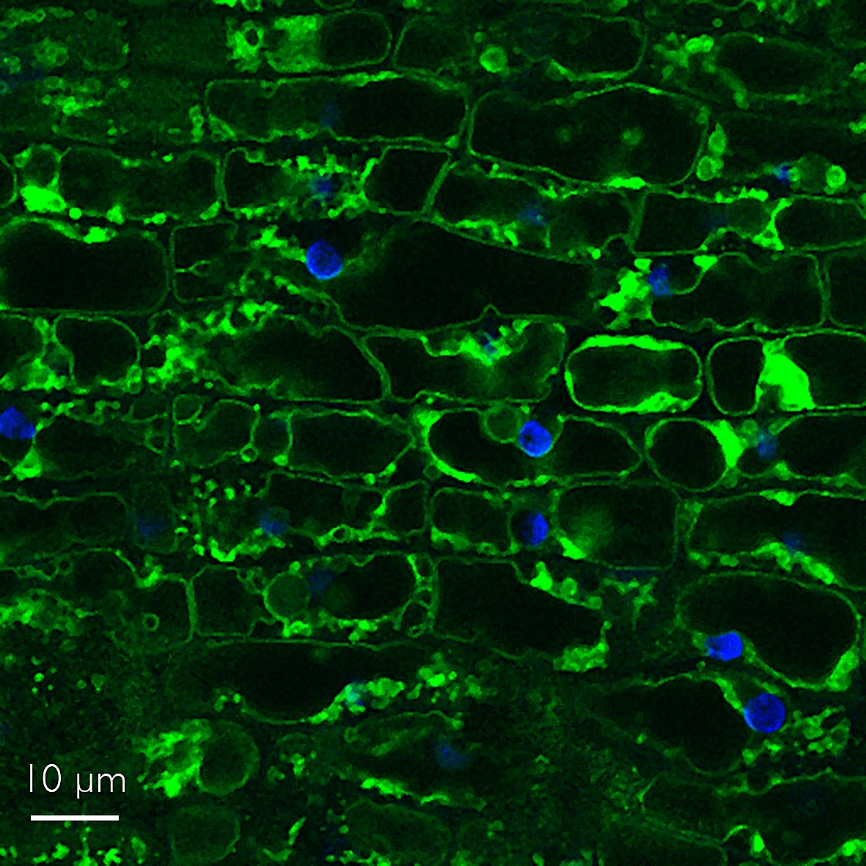

**Figure S6.**

Subcellular localization of *VviHT6* proteins to the tonoplast after stable expression in grapevine hairy roots.

The complete ORF of the *VviHT6* gene without the stop codon was amplified from Syrah cDNA (ripening berry) using the following primer pair

>HT6for CACCATGAACGGAGCTGTGCT

>HT6rev GTCATTCTTTGCTGCAGTAAC

before blunt end topo cloning, transfer to the pH7FWG2 vector and stable transformation of grapevine with agrobacterium rhyzogenes A4 according to Gomez et al., 2009.
